# C3 in the 10-20 system may not be the best target for the motor hand area

**DOI:** 10.1101/2023.02.06.527390

**Authors:** Hakjoo Kim, David L. Wright, Joohyun Rhee, Taewon Kim

**Author notes:** Corresponding author at: Washington University School of Medicine in St. Louis, 4444 Forest Park Avenue, St. Louis, MO 63108-2212, USA, E-mail address (Taewon Kim).

## Abstract

The C3 region in the international 10-20 system for electroencephalography (EEG) recording is assumed to represent the right motor hand area. Therefore, in the absence of transcranial magnetic stimulation (TMS) or a neuronavigational system, neuromodulation methods, such as transcranial direct current stimulation, target C3 or C4, based on the international 10-20 system, to influence the cortical excitability of the right and left hand, respectively. The purpose of this study is to compare the peak-to-peak motor evoked potential (MEP) amplitudes of the right first dorsal interosseus (FDI) muscle after single-pulse TMS at C3, C3h, and C1. Using an intensity of 110% of the resting motor threshold, 15 individual MEPs from each of C3, C3h, C1, and hotspots were randomly recorded from FDI for sixteen right-handed undergraduate students. Average MEPs were greatest at C3h and C1, with both being larger than those recorded at C3. These data are congruent with recent findings using topographic analysis of individual MRIs that revealed poor correspondence between C3/C4 and the respective hand knob. Implications for the use of scalp locations determined using the 10-20 system for localizing the hand area are highlighted.

**Highlights:**

- MEPs recorded at C3h and C1 were larger than those recorded at C3.
- Scalp locations other than C3 offer a more accurate estimation of human hand area.

## 1. Introduction

The number of studies using transcranial electrical stimulation, such as transcranial direct current stimulation (tDCS) and transcranial alternating current stimulation as a method of neuromodulation, continues to increase because it is generally inexpensive, simple, safe, and non-invasive (Buch et al., 2017). In a polarity-specific manner, tDCS has been shown to modify cortical excitability at the primary motor cortex (M1) (e.g., Nitsche and Paulus, 2000). Combining tDCS with physical practice has been reported to influence skill acquisition and/or retention (Reis et al., 2009; Buch et al., 2017). Determination of precise and reliable electrode placement for the application of tDCS, designed to target M1, often relies on the identification of the ‘motor hotspot’ using single-pulse transcranial magnetic stimulation (TMS) to elicit motor evoked potential (MEP) at the effector target site (e.g., first dorsal interosseous (FDI) muscle) (Groppa et al., 2012). In the absence of TMS, the placement of the electrode montage when administering tDCS often relies on a conventional positioning method, the international 10-20 system for electroencephalography (EEG) recording. In this case, the anode is located over the C3 or C4 region on the scalp when attempting to increase excitability of the right- or left-hand area of the M1, respectively (Brunoni et al., 2012; Silva et al., 2021). The selection of C3 or C4 is predicated on the assumption that this scalp location corresponds with the right- or left-hand area, often referred to as the ‘hand knob’ on the surface of M1 (Yousry et al., 1997).

Questions regarding C3/C4 as reliable estimates of the left- and right-hand knob have surfaced. Sparing et al. (2008) directly compared the accuracy of five unique methods for localizing the hand area of M1 prior to the application of TMS. Three methods took advantage of neuronavigation in conjunction with an individual’s structural- or functional-MRI data to facilitate positioning of the TMS coil on the scalp. Two conventional methods that relied only on standard cranial landmarks to locate the target area (i.e., C3 to target left hand knob) were also assessed. An M1 map was constructed using TMS to identify an MEP center of gravity (CoG). Sparing et al. argued that the MEP-CoG for a group of MEPs serves as a more accurate estimate of the scalp location directly above the region of maximum excitability (i.e., hotspot) as opposed to just using the location in the grid that is associated with the largest MEP. Each individual’s MEP-CoG was used as a reference for the location of the hand knob. The international 10-20 system, one of the conventional approaches evaluated, exhibited the poorest correspondence with the hand area as C3 was associated with the largest spatial distance from the motor map’s CoG. In addition, the smallest amplitude TMS-elicited MEPs normalized to the motor map CoG (i.e., ~49%) were recorded at C3.

More recently, Silva et al. (2021) used topographic analysis of individual MRIs to evaluate and compare the accuracy of C1/C2, C3h/C4h, and C3/C4 points for locating the M1 hand areas on the scalp. C1/C2 are additional sites derived from the 10-20 system, whereas C3h or C4h are scalp locations between C1/C3 and C2/C4, respectively, derived from use of the international 10-5 system. The hand knob was identified from anatomical landmarks for 30 individuals’ MRIs. Specifically, the hook sign in the sagittal plane and an inverted omega or horizontal epsilon sign in the horizontal plane of the structural MRI was used to determine the location of the hand area (see Yousry et al., 1997; Silva et al., 2021). The distance between C1/C2, C3h/C4h, and C3/C4 from the individual’s hand area was subsequently determined. Congruent with the findings of Sparing et al. (2008), C3 and C4 again revealed the largest deviation from the hand knob. Moreover, the intermediate scalp locations, namely C3h and C4h derived from the 10-5 system, offered the most accurate approximation of the location of the hand area.

The purpose of this study was to extend the work of Silva and colleagues by comparing the peak-to-peak amplitude of the MEP from the FDI muscle of the right hand from TMS administered at scalp locations used by Silva et al. (2021), namely, C3, C3h, and C1. Based on the work of Silva et al., we anticipate that TMS-elicited MEPs from C3h should be larger than those obtained at C3. Because Silva et al.’s work also reported that C1 exhibited a smaller spatial deviation from the hand knob than C3, we expect relatively larger MEPs at C1 when compared to C3.

## 2. Results

Mean peak-to-peak amplitude of TMS-elicited MEPs for each individual recorded from C3, C3h, and C1 were submitted to a repeated-measures analysis of variance (ANOVA) (Region: C3, C3h, C1). Mauchly’s test revealed that the assumption of sphericity was violated (*p* = .039), leading to a Greenhouse-Geisser correction in degrees of freedom. The ANOVA revealed a significant main effect of Region, *F*(1.458, 21.870) = 4.956, *p* = .025. Post hoc analysis indicated that this main effect was a result of the mean peak-to-peak amplitude of TMS-elicited MEP from C3 (*M* = 0.063, *SD* = 0.149) being significantly lower than the mean peak-to-peak amplitude of TMS-elicited MEP from C3h (*M* = 0.661, *SD* = 0.803, *p* = .020) and C1 (*M* = 0.856, *SD* = 0.959, *p* = .019). Peak-to-peak amplitude of TMS-elicited MEP recorded from C3h and C1 did not differ significantly (see Figure 1).

**Fig. 1.**
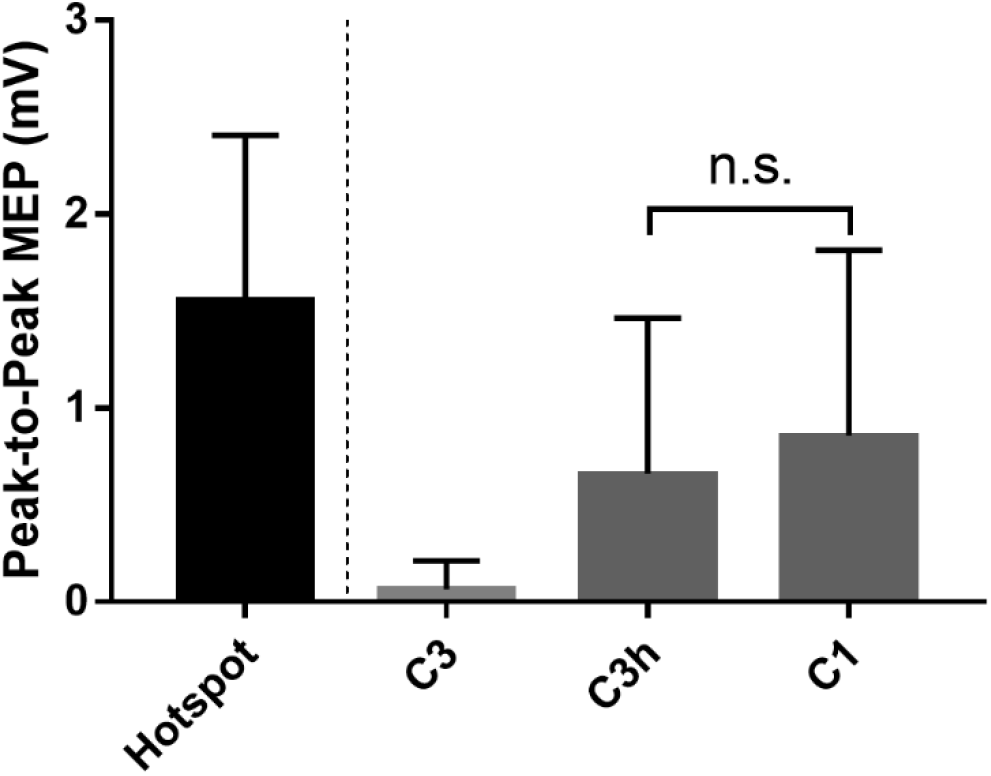
The mean peak-to-peak MEPs at the right FDI muscle from four TMS targets (FDI hotspot, C3, C3h, and C1). The error bars represent the standard deviation.

## 3. Discussion

The mean peak-to-peak amplitude of TMS-elicited MEPs was largest at scalp locations C3h and C1, with both being greater than those recorded at C3 (Figure 1). Since lower MEPs are expected when stimulation is further from the hand area, these data are congruent with recent concerns about the efficacy of C3 or C4 as accurate estimates of this region on the M1 (Sparing et al., 2008). The present data are also in keeping with recent findings from Silva et al. (2021) that used topographic analysis to reveal closer spatial proximity of the C3h and C4h scalp locations to the respective hand knobs on the M1 than C3 and C4. While the mean peak-to-peak amplitude of TMS-elicited MEP at C1 was greater than that observed at C3h (see Figure 1), the difference was not significant. This finding was not totally unexpected given Silva et al. reported that the spatial deviation of C3h/C4h (~0.98 cm) and C1/C2 (~ 1.18 cm) from the hand knob did not differ significantly.

The poor correspondence between C3/C4 and the hand area in the present work is consistent with earlier evidence from Sparing et al. (2008). This work noted that targeting M1 using the 10-20 system was the least accurate of five distinct approaches, which included at least one other method that relied only on cranial landmarks to identify the scalp location assumed to overlay the hand area. These data provide further evidence suggesting that more accurate localization of the hand knob at M1 can be accomplished by utilizing scalp sites derived from the from the 10-5 system or an alternative location (i.e., C1) from the more commonly used 10-20 system (see also Herwig et al., 2003). The lack of difference observed in the mean peak-to-peak amplitude of TMS-elicited MEPs was largest at scalp locations C3h and C1 is consistent with the claim that a scalp position between C1-C3h and C2-C4h may more precisely locate the target hand area (Silva et al., 2021).

The present data and those of Silva and colleagues have some important practical implications given many current day clinical practice and research efforts routinely use C3/C4 to target the hand area (see also Brunoni et al., 2012). Importantly, these data highlight that localizing the target region at M1 can be improved using alternative scalp sites that are easily located and do not necessitate the use of neural navigation. Clearly, the use of a conventional TMS protocol to determine the motor hotspot prior to conducting neuromodulation using transcranial electrical stimulation tools such as tDCS is preferable. Indeed, it would be easy to assume that most clinical and research settings engaging in modifying cortical excitability via tDCS are equipped with this capability. However, a brief review of the published articles indexed by PubMed for the year 2022^1^ revealed that 67.6% of papers in which the hand area of M1 was targeted relied on the international 10-20 system for placement of the electrodes. Moreover, on the clinical front, there is increasing interest in the therapeutic use of non-invasive brain stimulation tools, with considerable focus on tDCS because of the ease of administration. The consensus is that increasing the length of tDCS after-effects will be critical for functional efficacy which will likely demand the administration of repeated doses (Brunoni et al., 2012). This requirement will intensify the need to provide this form of treatment in the home setting to improve access and reduce the various burdens placed on patients and their caregivers when traveling to clinics and hospitals (Charvet et al., 2020). Clearly, in these circumstances, localization for electrode placement will almost certainly be dependent on the use of conventional positioning methods that use cranial landmarks only. Evidence presented herein and elsewhere suggests that successful targeting of the hand area in these situations, as well as more routine experimental settings, can be accomplished using C3h in the 10-5 system or C1 rather than C3 from the 10-20 system.

### 3.1. Limitations

While a single experienced experimenter identified scalp locations in accordance with international 10-20 and 10-5 systems to reduce measurement error, measurement error may have occurred when locating C3, C3h, and C1 (e.g., uncertain inion). In addition, errors may have also occurred when TMS was administered at each scalp location (i.e., C3, C3h, and C1). The magnitude of error for positioning the TMS coil in x, y, and z planes was recorded during each stimulation, and mean errors were formally assessed for each scalp location (see Table 1).

**Table 1.** Mean error information at each TMS target. The information involves target (mm), angular (degree), and twist errors (degree).

| Target | Target Error (*SD*) | Angular Error (*SD*) | Twist Error (*SD*) |
| --- | --- | --- | --- |
| Hotspot | 0.39 (0.23) | 0.22 (0.12) | 0.03 (0.10) |
| C1 | 0.36 (0.22) | 0.25 (0.13) | 0.04 (0.11) |
| C3h | 0.40 (0.21) | 0.24 (0.15) | 0.05 (0.11) |
| C3 | 0.45 (0.27) | 0.26 (0.15) | 0.06 (0.14) |
| Mean | 0.40 (0.23) | 0.24 (0.14) | 0.05 (0.11) |

## 4. Experimental Procedure

### 4.1. Participants

Twenty-three right-handed undergraduate students participated in this study. All participants provided a TMS screening form and written informed consent before participating in this experiment, which was approved by the Texas A&M University Institutional Review Board. Additionally, individuals completed the Edinburgh Handedness Inventory (Oldfield, 1971) prior to the experiment, and those who were identified as left-handed or ambidextrous were excluded (two left-handed; four ambidextrous). In addition, one subject was excluded from the study because the head size could not be correctly measured due to the hairstyle (i.e., dreadlocks). Thus, only data from sixteen right-handed undergraduate students (thirteen females and three males; mean age: 19.69 (*SD*: 0.87)) were included in the data analyses.

### 4.2. Transcranial magnetic stimulation (TMS)

For MEP measurement, single-pulse TMS was administered by DuoMAG MP-Dual (DEYMED Diagnostic s.r.o., Czech Republic) with a figure-of-eight butterfly coil (2 × 70 mm-diameter windings, DUOMAG 70BF COIL, DEYMED Diagnostic s.r.o., Czech Republic). In addition, the Brainsight TMS Navigation system (Version 2.4.3, Rogue Research, Canada) was utilized for navigated-TMS, enabling sustained stimulation of the same target points. A hotspot of the FDI muscle was determined based on the peak-to-peak MEP amplitudes obtained from pre-set-grid-based hotspot-hunting (grid spacing: 2 by 2 mm). The hotspot-hunting was performed with a coil orientation of a 45-degree angle. After finding a hotspot in each subject, the resting motor threshold (rMT) of the FDI muscle was determined as a minimum TMS intensity to evoke the peak-to-peak amplitude of 0.05 mV in at least 5 out of 10 trials (mean rMT: 39.50 (*SD*: 6.94)). Single-pulse TMS at an intensity of 110% of rMT (see Kallioniemi and Julkunen, 2016) was administered at C3, C3h, C1, and a hotspot fifteen times each in random order (e.g., fifteen pulses at C1, a hotspot, C3, and C3h, respectively). Single-pulse TMS was given irregularly so that participants could not predict the timing of the stimulation, and the time interval between each TMS pulse was about 5 seconds. The target TMS coil orientation was maintained 45 degrees to the midline of the brain (see Table 1).

### 4.3. Electromyography (EMG)

Navigated-TMS was delivered at C3, C3h, C1, and a pre-determined hotspot for FDI to induce an MEP for the right FDI muscle. EMG signals were recorded through disposable Ag-AgCl electrodes (10-mm diameter, Adult Tape ECG Cleartrace, CONMED, US), amplified by NL844 AC Pre-amplifier (Gain ×100, Digitimer Ltd., UK), which was connected to NL820A Isolation Amplifier (Gain ×1, Digitimer Ltd., UK), filtered by NL136 Four Channel Low Pass Filters (2 kHz, Digitimer Ltd., UK), and sampled at 5 kHz by Signal software (Version 7.04, CED Ltd., UK). TMS-induced MEPs were recorded from the right FDI muscle, and the background noise was kept to less than 0.03 mV.

### 4.4. Procedure

The experiment was performed on a single day, and all participants completed a consent form and pre-experiment questionnaires prior to participation. First, an experimenter measured the head size of the participants to determine each target point (i.e., C3, C3h, and C1). Then, two EMG electrodes were attached to the right hand (FDI muscle). A tracker for the neuronavigation system was attached to the forehead after confirming background noise and signal response to participants’ finger movements. A co-registration procedure was performed using a template head (i.e., the MNI ICBM 152 average brain), after which an FDI hotspot and rMT of the FDI muscle were determined. Using single-pulse TMS at an intensity of 110% of the FDI rMT, the peak-to-peak MEPs from C3, C3h, and C1, as well as from an FDI hotspot, were measured fifteen times each in random order.

### 4.5. Data analysis

The primary dependent variable was the peak-to-peak MEPs of the FDI muscle. The peak-to-peak MEPs were captured by Signal (Version 7.04, CED Ltd., UK). Statistical analysis was performed using SPSS (Version 28, IBM, US).

## Acknowledgments

The authors would like to thank Drs. John Buchanan and Yuming Lei for their feedback on this study. We also thank the reviewers for their valuable comments.

## Abbreviations

ANOVA: analysis of variance
EEG: electroencephalography
EMG: electromyography
FDI: first dorsal interosseus
M: mean
MEP: motor evoked potential
rMT: resting motor threshold
SD: standard deviation
tDCS: transcranial direct current stimulation
TMS: transcranial magnetic stimulation

## Footnotes

1 The key phrase “tDCS” and “M1” were used to search all articles listed in PubMed (https://pubmed.ncbi.nlm.nih.gov/) during the period January 1, 2022-December 31, 2022.

